# Urinary exfoliated cells exhibit mesodermal lineage differentiation potential in the absence of dedicated culture conditions

**DOI:** 10.1101/2025.08.31.673107

**Authors:** Naonori Kumagai, Takuma Ando, Tomomi Kondoh, Yuji Matsumoto, Michiaki Abe, Yohei Ikezumi

## Abstract

**Introduction:** Urine contains various cells derived from the kidney, including renal tubular epithelial cells, glomerular epithelial cells, and mesenchymal stem cells. Urinary exfoliated mesenchymal stem cells can be differentiated into adipocytes, osteoblasts, chondrocytes, muscle cells, or neuronal cells, if grown in specialized culture conditions. In this study, however, we identified the development of multiple mesodermal cell lineages from urinary exfoliated cells despite using a single culture system not designed for these cell differentiations.

**Methods:** Urinary exfoliated cells used for culturing were obtained from pediatric patients with kidney disease. All cultures were maintained using an identical, non-specifically supplemented, rich, DMEM/F12-based growth medium, refreshed regularly throughout the experiments. Cells of different mesodermal lineages—adipocytes, osteoblasts, and chondrocytes—were identified by staining with Oil Red O, Alizarin Red, and Alcian Blue solutions. Moreover, the expression of mRNA markers specific for adipocytes, osteoblasts, and chondrocytes, as well as for stem cells and—more specifically—mesenchymal stem cells, was analyzed by RT-PCR.

**Results:** Although the abundances varied, in all 11 cultures of urinary exfoliated cells from six patients, dye staining revealed multiple cell clusters of mesodermal lineages. All cultures contained clusters of the adipocyte and chondrocyte lineages, and seven also contained clusters of the osteoblast lineage. Morphology differences were found even between clusters stained with the same dye. RT-PCR confirmed the presence of the three lineages, while also revealing expression of genes specific to pluripotent stem cells (*Nanog, Oct3/4, SOX2*, and *LIF*) and genes more specific for mesenchymal stem cells (*CD73, CD90*, and *CD105*).

**Conclusions:** Among urinary exfoliated cells cultured using our method, clusters of cells of mesodermal lineages were observed without needing to selectively adjust the culture medium. The cells seemed to differentiate and proliferate from mesenchymal stem cells in the urine. We envision that these initial findings may eventually contribute to a deeper understanding of urinary stem cell differentiation and to optimizing lineage-specific differentiation for research and therapeutic applications.

## 1. Introduction

Stem cells possess self-renewal capacity and multipotent differentiation potential [1]. Mesenchymal stem cells can differentiate into various mesodermal cell lineages, including adipocytes, osteoblasts, chondrocytes, muscle cells, and neuronal cells. In cell culture, such differentiations are usually achieved by specifically adjusted conditions. Mesenchymal stem cells have shown potential as a source for cell therapy in a wide range of conditions—including burns, wound healing, and neurological, cardiac, respiratory, metabolic, endocrine, bone, and renal diseases [2]. Regarding the latter, applications have been explored for acute kidney injury, chronic kidney disease, diabetic nephropathy, lupus nephritis (LN), and kidney transplantation [3]. Differentiation ability and morphology of mesenchymal stem cells vary depending on the tissue source and donor age [2].

Urine contains live cells derived from the renal tissue, such as renal tubular epithelial (RTE) cells and glomerular epithelial cells, which can be cultured [4][5][6]. Urine also contains mesenchymal stem cells thought to be derived from the kidney, which can differentiate into various mesodermal lineage cells under specialized culture conditions [7][8]. Regarding clinical purposes, these urinary mesenchymal stem cells show similar potential as other mesenchymal stem cells [7].

We developed a unique method to culture urinary exfoliated cells, which we have primarily used to study RTE cells [9][10]. These cultured RTE cells have been investigated for transporter functions [9], proliferation ability [11], mRNA expression [12], and 3D tubule reconstruction [10]. However, we have not previously examined whether this method can also support the growth or differentiation of other cell types.

In the present study, we applied our culture method to urinary exfoliated cells from pediatric patients and identified multiple cell types with staining and gene expression profiles consistent with the development of distinct mesodermal lineages, including adipocytes, osteoblasts, and chondrocytes. The presence of stem-cell-associated and mesenchymal markers suggests that these differentiated cell types arose from mesenchymal stem-like cells present in the urine, even without the use of specialized differentiation protocols.

## 2. Methods

### 2.1 Donors and Cell Culture

Urinary exfoliated cells were cultured using our previously established method [9][11]. Samples were obtained from pediatric patients with kidney disease, diagnosed at and admitted to the Department of Pediatrics at Fujita Health University Hospital, Japan. In brief, 50–200 mL of voided urine was collected from each patient using a clean-catch procedure. The urine samples were centrifuged at 280 g for 10 min at room temperature, and the pellets were washed twice at 280 g for 10 min with minimum essential medium (MEM) (Gibco, MA, USA) containing 20% fetal bovine serum (FBS) (Gibco), 100 U/mL penicillin, and 100 μg/mL streptomycin (Gibco). Cells were then washed once with the growth medium, after which they were resuspended in the growth medium and seeded into the multiple wells of a 6-well culture plate with a diameter of 3.5 cm each (AGC techno glass, Shizuoka, Japan) coated with Cellmatrix Type 1-C collagen (Nitta Gelatin, Osaka, Japan). The growth medium consisted of Dulbecco’s modified Eagle’s medium/Ham’s F12 (DMEM/F12) (Wako, Osaka, Japan), 10% FBS, 5 μg/mL insulin, 5 μg/mL transferrin, 5 ng/mL sodium selenite (Sigma-Aldrich, MO, USA), 10^-8^ M dexamethasone (Sigma-Aldrich), 10^-8^ M T3 (Sigma-Aldrich), and 5 mM nicotinamide (Wako). Cultures were maintained at 37 °C and 5% CO_2_. The growth medium was first replaced 10–14 days after seeding and then every 3–4 days. After reaching confluency or cell proliferation arrest, cells were cultured for an additional 7–14 days.

### 2.2 Specific Cell Staining

Whole-well cell cultures were subjected to lineage-specific staining procedures. After aspirating the medium, the cells were fixed with 4% paraformaldehyde (Wako) at room temperature for 20 min and washed three times with phosphate-buffered saline (PBS; Wako) at room temperature for 5 min. Adipocytes were stained using Oil Red O solution (Muto Pure Chemicals, Tokyo, Japan). Osteoblasts were stained using 1% Alizarin Red solution pH 6.3–6.4 (Muto Pure Chemicals). Chondrocytes were stained using Alcian Blue solution pH 2.5 (Muto Pure Chemicals). Stained cells were observed and photographed using a BZ-x810 fluorescence microscope (KEYENCE, Osaka, Japan) and analyzed using BZ-X800 Analyzer software (KEYENCE).

### 2.3 RT-PCR

Total RNA was extracted per well of cultured cells using RNeasy (QIAGEN, Hilden, Germany) according to the manufacturer’s instructions. cDNA was synthesized from 2 μg of RNA using SuperScript III First-Strand Synthesis SuperMix (Invitrogen, CA, USA) according to the manufacturer’s protocol. PCR was performed in a 25-μL reaction system using Taq DNA polymerase (Takara Bio, Shiga, Japan) with 0.5 μL of cDNA per reaction. For primers see Table 1. Cycling conditions were as follows: denaturation at 95 □ for 1 min, annealing at 55 □ for 1 min, elongation at 72 □ for 1 min, and a final extension at 72 □ for 5 min. Amplified products (8 μL) were electrophoresed using 1.2%–3% agarose gels (Agarose-S; NIPPON GENE, Tokyo, Japan) containing Midori Green Advance (NIPPON Genetics, Tokyo, Japan) and imaged using a WSE-5400 Printgraph Classic UV transilluminator (ATTO, Tokyo, Japan).

**Table 1.**
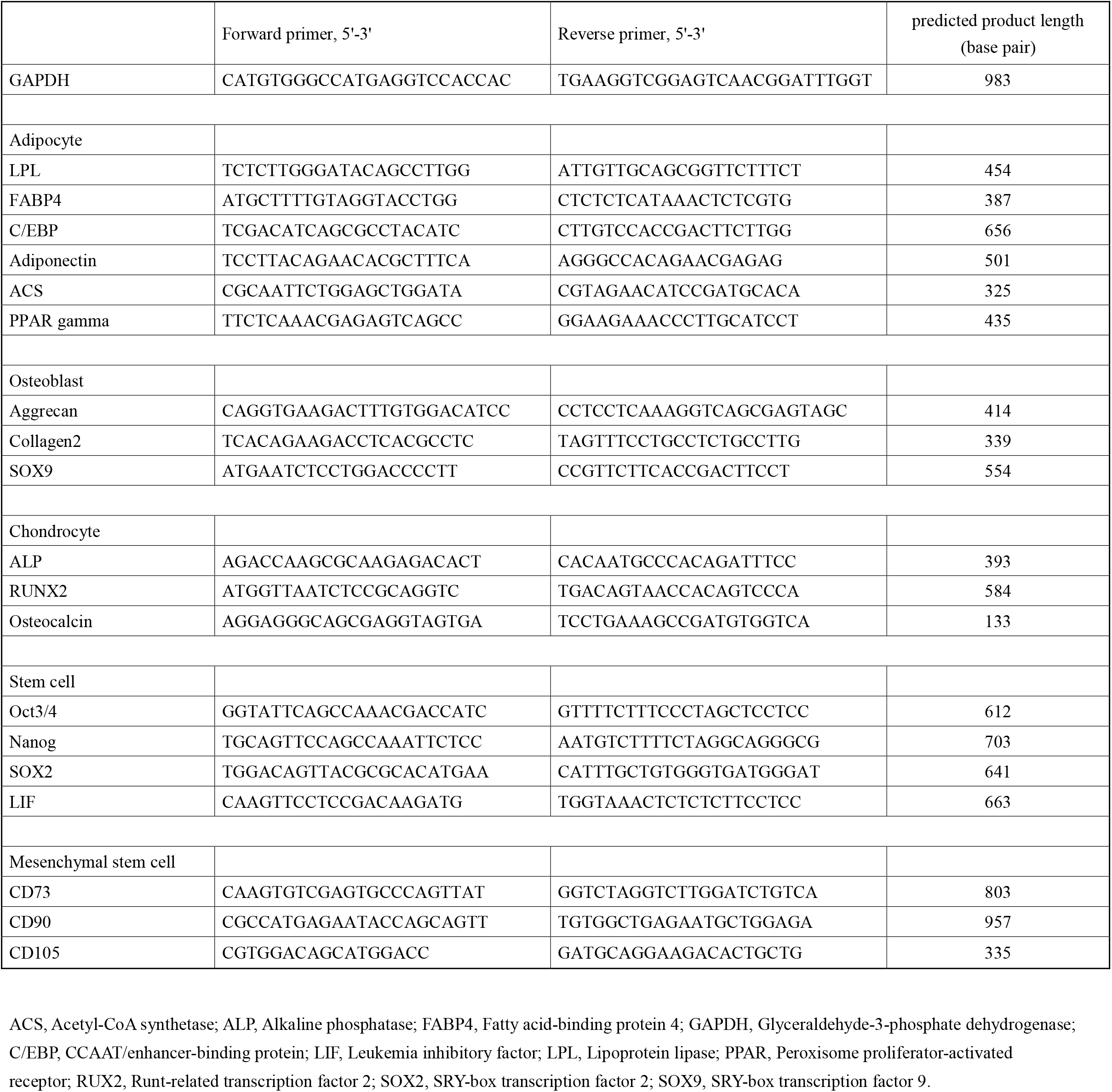
PCR primers used in the experiments.

### 2.4 Ethics

This study was conducted with the approval of the Fujita University Ethics Committee (HM20-32) and in accordance with the 1964 Declaration of Helsinki.

## 3. Results

### 3.1 Subjects

Pediatric patients with various kidney diseases participated in this study. One or more urine samples were collected from each patient, and each sample was used to establish a separate cell culture (Table 2).

**Table 2.** Demographics of the patients and the cell staining results.

| Patient | Sex | Age<br>(yr) | Diagnosis | Creatinine<br>(mg/dL) | U-pro/Cr<br>(g/g) | U-RBC<br>(/HPF) | Oil Red O | %PSA | Alizarin Red | %PSA | Alcian Blue | %PSA |
| --- | --- | --- | --- | --- | --- | --- | --- | --- | --- | --- | --- | --- |
| Patient 1 | M | 4 | HSPN |  |  |  |  |  |  |  |  |  |
| Culture 1 |  |  |  | 0.34 | 3.1 | 100< | + | 4.51 | + | 6.40 | + | 4.91 |
| Culture 2 |  |  |  | 0.33 | 1.01 | 100< | + | 6.60 | + | 1.62 | + | 12.40 |
| Culture 3 |  |  |  | 0.29 | 0.12 | 50-99 | + | 9.48 | + | 0.01 | + | 4.91 |
| Patient 2 | M | 15 | IgAN |  |  |  |  |  |  |  |  |  |
| Culture 4 |  |  |  | 0.75 | 1.25 | 50-99 | + | 11.36 | + | 0.05 | + | 7.79 |
| Patient 3 | F | 14 | LN |  |  |  |  |  |  |  |  |  |
| Culture 5 |  |  |  | 0.65 | 4.02 | 50-99 | + | 1.89 | - | 0 | + | 1.70 |
| Culture 6 |  |  |  | 0.65 | 1.42 | 50-99 | + | 1.18 | + | 0.02 | + | 16.91 |
| Patient 4 | F | 14 | ANCA |  |  |  |  |  |  |  |  |  |
| Culture 7 |  |  |  | 1.14 | 0.455 | 100< | + | 17.03 | + | 4.84 | + | 2.59 |
| Culture 8 |  |  |  | 0.96 | 0.323 | 30-49 | + | 6.44 | + | 9.04 | + | 1.57 |
| Patient 5 | F | 12 | MN |  |  |  |  |  |  |  |  |  |
| Culture 9 |  |  |  | 0.36 | 20.33 | - | + | 1.24 | - | 0 | + | 2.52 |
| Culture 10 |  |  |  | 0.36 | 3.98 | - | + | 2.52 | - | 0 | + | 4.29 |
| Patient 6 | M | 11 | HSPN |  |  |  |  |  |  |  |  |  |
| Culture 11 |  |  |  | 0.52 | 2.71 | 50-99 | + | 0.01 | - | 0 | + | 2.54 |

ANCA, anti-neutrophil cytoplasmic antibody associated glomerulonephritis; HPF, high power field; n.d., not done; HSPN, Henoch-Schönlein purpura nephritis; IgAN, IgA nephropathy; LN, lupus nephritis; MN, membranous nephropathy; %PSA, percentage of the positive stained area per well; u-pro/cr, urinary protein/creatinine; u-RBC, urinary red blood cell.

### 3.2 Identification of Adipocytes, Osteoblasts, and Chondrocytes by Cell Staining

Cell clusters positive for Oil Red O, Alizarin Red, and Alcian Blue staining were detected, suggesting the presence of adipocytes, osteoblasts, and chondrocytes, respectively (Fig. 1). The staining patterns of these clusters varied, and cell morphology could differ even between clusters positive for the same dye (Fig. 1, Table 2). The percentage of the positive stained area per well (%PSA values) differed per culture, from barely present (0.01%) to 17% (Table 2). All cultures were positive for Oil Red O and Alcian Blue staining, but only seven of the 11 cultures included Alizarin Red-positive cell clusters (Table 2).

**Figure 1.**
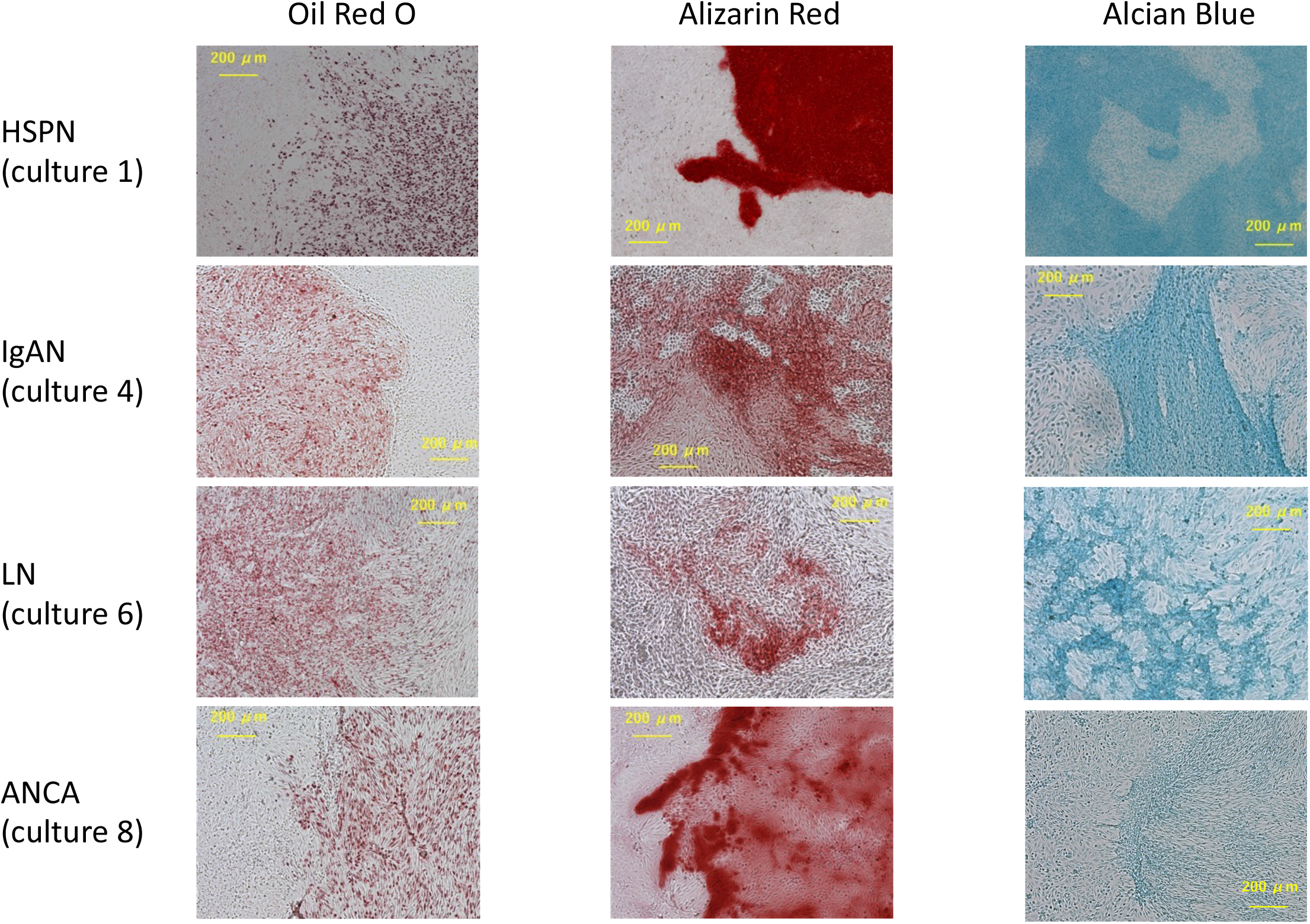
Staining with lineage-specific dyes of cultured urinary exfoliated cells Mesodermal lineages identified were adipocytes (by Oil Red O solution staining), osteoblasts (by Alizarin Red staining), and chondrocytes (by Alcian Blue staining). ANCA, anti-neutrophil cytoplasmic antibody associated glomerulonephritis; HSPN, Henoch-Schönlein purpura nephritis; IgAN, IgA nephropathy; LN, lupus nephritis.

As an example of unexplained differences, for the two patients with Henoch-Schönlein purpura nephritis (HSPN; Patients 1 and 6), the staining results differed. Patient 1 exhibited clusters for all three lineages, whereas only adipocyte-like and chondrocyte-like clusters were observed for Patient 6 (Table 2).

Differences in abundances of each type of cluster were also noted between cultures derived from separate urine samples from the same patient (Table 2).

### 3.3 RT-PCR Analysis of Gene Expression Characteristic for Adipocytes, Osteoblasts, and Chondrocytes

Cell cultures Nos. 1, 4, 6, and 8—which stained positive for all three mesodermal cell types—were analyzed by RT-PCR for transcription of lineage-specific markers of adipocytes, osteoblasts, and chondrocytes [8]. Marker expression profiles for each of the three cell types varied between cultures, and the combination of positive and negative findings suggests the presence of cells in partially differentiated states (Fig. 2).

**Figure 2.**
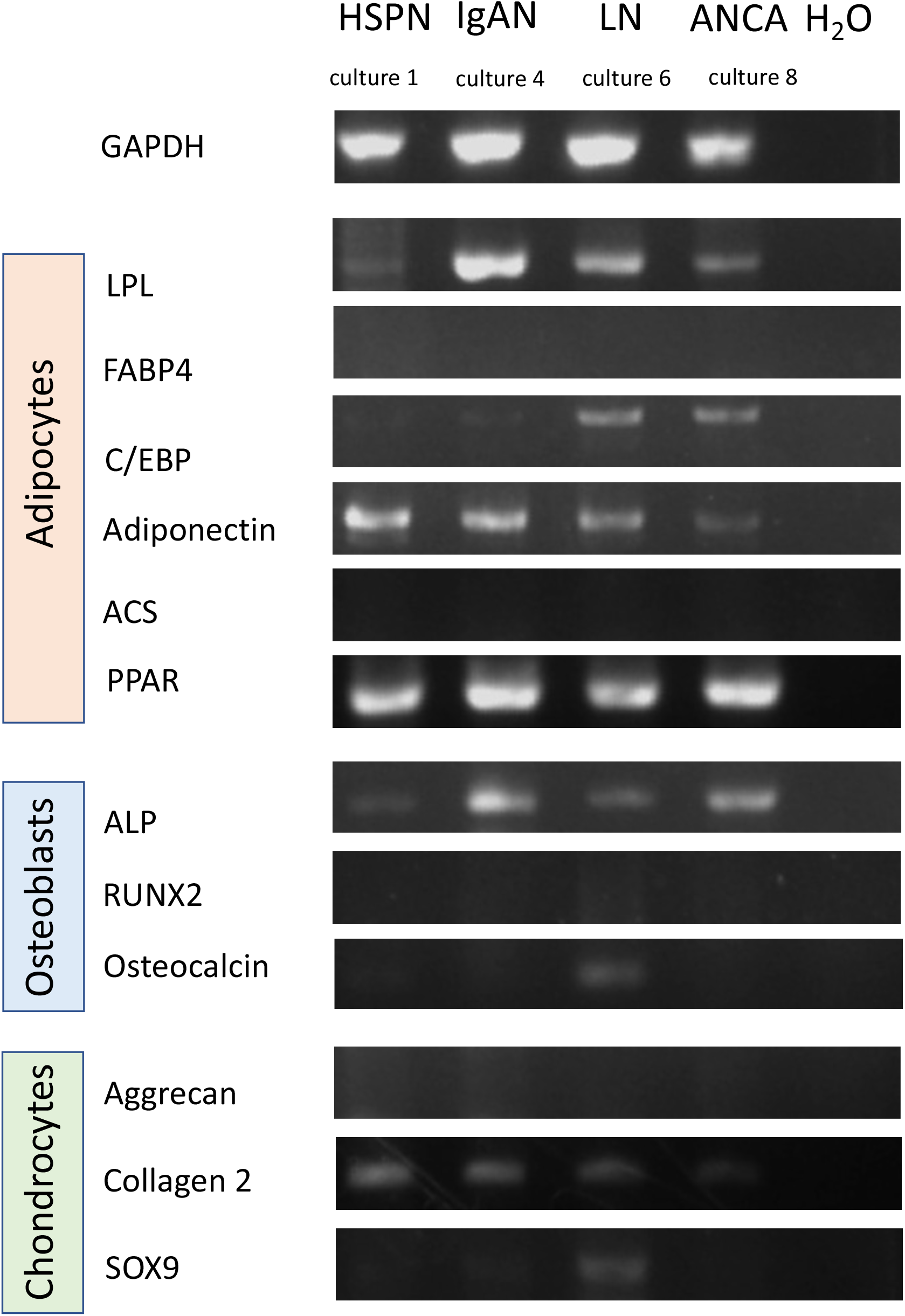
Detection of specific gene marker expression for adipocytes, osteoblasts, and chondrocytes by RT-PCR ACS, Acetyl-CoA synthetase; ALP, Alkaline phosphatase; ANCA, anti-neutrophil cytoplasmic antibody associated glomerulonephritis; C/EBP, CCAAT/enhancer-binding protein; GAPDH, Glyceraldehyde-3-phosphate dehydrogenase; FABP4, Fatty acid-binding protein 4; HSPN, Henoch-Schönlein purpura nephritis; IgAN, IgA nephropathy; LN, lupus nephritis; LPL; Lipoprotein lipase; PPAR, Peroxisome proliferator-activated receptor; RUX2, Runt-related transcription factor 2; SOX9, SRY-box transcription factor 9.

### 3.4 RT-PCR analysis of gene expression characteristic for stem cells

The same RNA samples were also investigated by RT-PCR for stem cell-associated transcripts. All four cultures expressed the pluripotency-associated markers *Nanog, Oct3/4*, and *LIF* [1][13], with *SOX2* expressed at trace levels (Fig. 3). Additionally, all cultures expressed *CD73, CD90*, and *CD105*, which are characteristic markers of mesenchymal stem cells [2] (Fig. 4).

**Figure 3.**
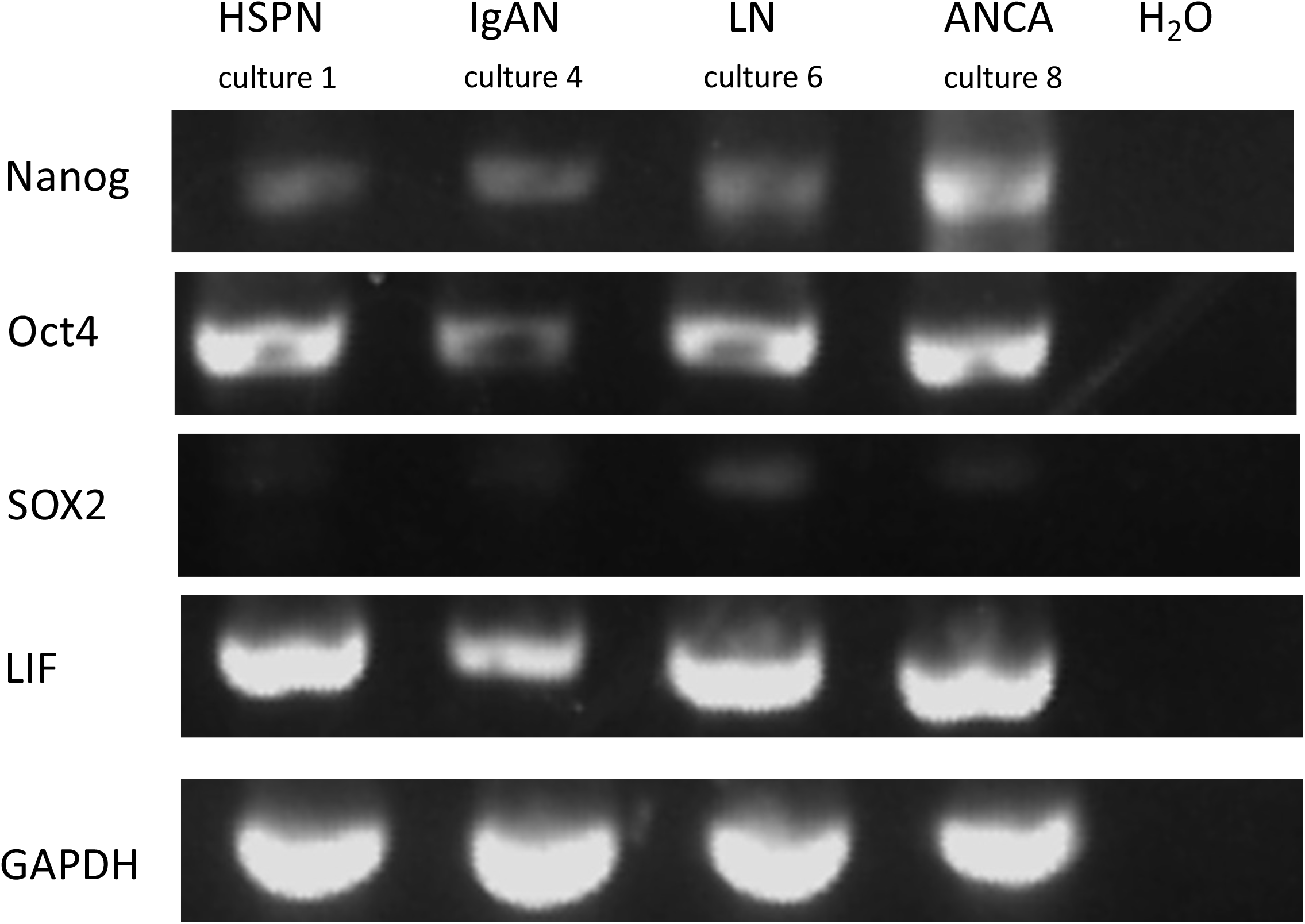
Detection of stem cell gene marker expression by RT-PCR ANCA, anti-neutrophil cytoplasmic antibody associated glomerulonephritis; GAPDH, Glyceraldehyde-3-phosphate dehydrogenase; HSPN, Henoch-Schönlein purpura nephritis; IgAN, IgA nephropathy; LIF, Leukemia inhibitory factor; LN, lupus nephritis; SOX2, SRY-box transcription factor 2.

**Figure 4.**
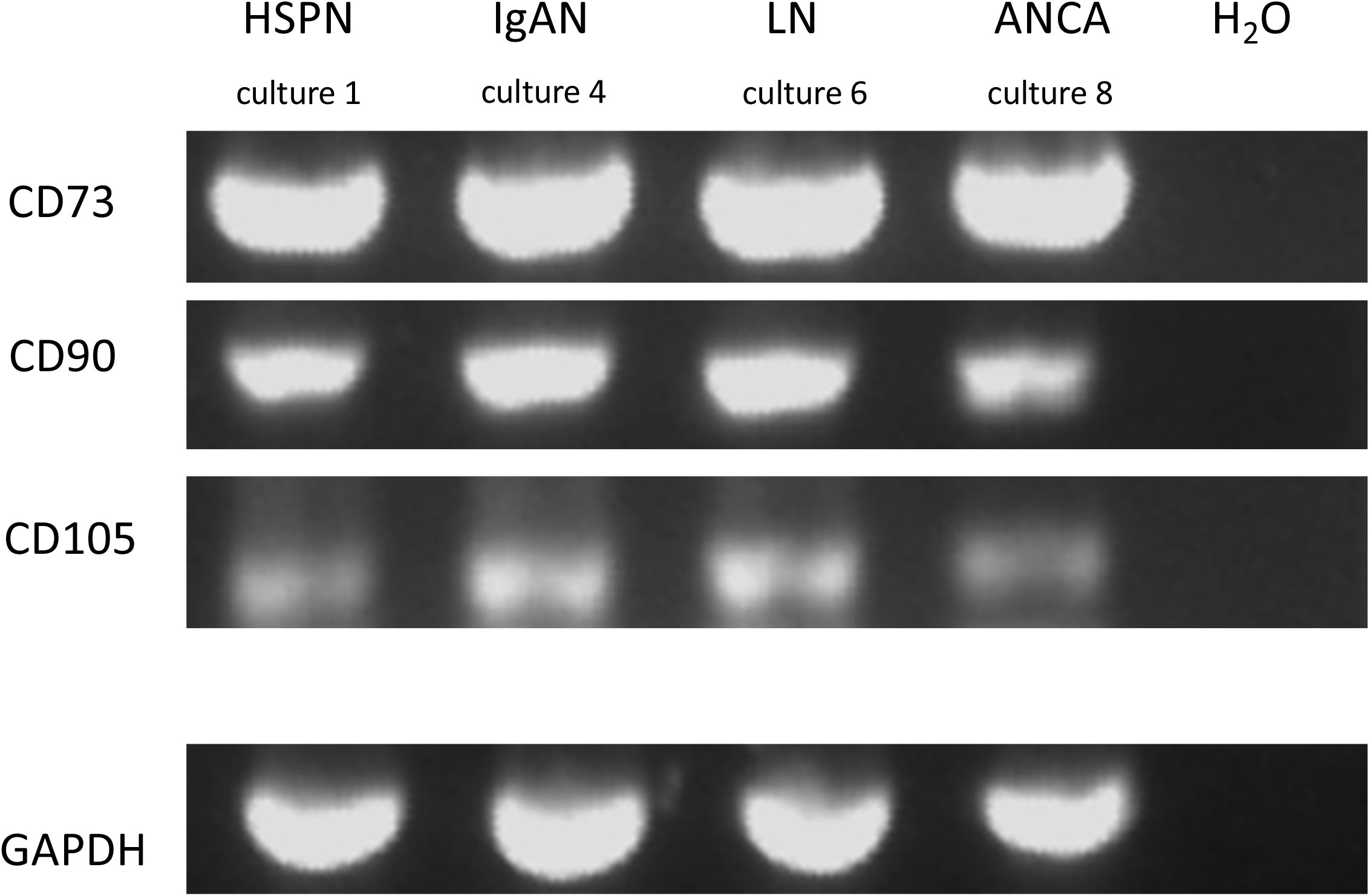
Detection of mesenchymal stem cell gene marker expression by RT-PCR ANCA, anti-neutrophil cytoplasmic antibody associated glomerulonephritis; GAPDH, Glyceraldehyde-3-phosphate dehydrogenase; HSPN, Henoch-Schönlein purpura nephritis; IgAN, IgA nephropathy; LN, lupus nephritis.

## 4. Discussion

This study demonstrates that urinary exfoliated cells from pediatric patients with kidney disease can give rise to adipocyte-, osteoblast-, and chondrocyte-like cells even when cultured in a single, non-specialized growth system. These three cell types are not normally present in urine and likely originated from kidney-derived mesenchymal stem cells shed into the urine. Although urinary mesenchymal stem cells have previously been shown to differentiate into multiple mesodermal lineages under tailored conditions [8], this is, to our knowledge, the first report of such differentiation occurring without specialized media.

Specific staining and RT-PCR confirmed the presence of adipocyte-, osteoblast-, and chondrocyte-like cells regardless of the donor’s kidney disease, indicating that this ability is not restricted to a particular pathology. However, not all three lineages appeared in every culture, even when derived from the same patient. Within individual cultures, multiple clusters of the same lineage were often present, and these could differ in morphology and staining pattern. Gene expression of lineage-specific markers, assessed at the culture level, also varied between cultures. These observations suggest heterogeneity in the differentiation state of the cells, likely reflecting differences in the initial urinary cell populations and chance events during culture establishment.

The detection of *Nanog, Oct3/4, SOX2, LIF, CD73, CD90*, and *CD105* expression in the cultured cells indicates the presence of mesenchymal stem cells among the urinary exfoliated cells, consistent with previous reports [8]. Future work that isolates these stem cells directly from urine or cultured exfoliated cells could help to clarify their differentiation potential.

Until now, it has been generally accepted that inducing the differentiation of stem cells into adipocytes, osteoblasts, and chondrocytes requires appropriate adjustments to the culture conditions (e.g., the medium contents): adipocytes mainly require isobutylmethylxanthine (IBMX), dexamethasone (DEX), insulin, and indomethacin [14]; osteoblasts require β-glycerophosphate, DEX, and ascorbate [15]; and chondrocytes require TGF-β, DEX, a mixture of insulin, transferrin, and selenium (ITS), ascorbate, and BMP [16]. The culture medium used in the present study lacked (relevant amounts of) some of these components—e.g., IBMX for adipocytes, β-glycerophosphate for osteoblasts, and TGF-β and BMP for chondrocytes—and therefore, differentiation into any of these three cell types was not expected.

The urinary stem cells obtained using our culture method exhibited pronounced differentiation abilities, as multiple mesodermal cell types developed even without specialized culture conditions. Possible reasons for this unexpected capability include: (1) the stem cells were derived from urine samples obtained from children, whose cells generally have a higher differentiation capacity than those from older individuals [9][17]; (2) the cells originated from patients with kidney disease, which may affect their characteristics, as culture efficiency has been reported to decrease when kidney disease activity subsides [9]; (3) the stem cells may undergo partial differentiation during their transit from the kidney to the urine and subsequently into culture; (4) some cells with feeder-like properties may be present in the cultured urinary exfoliated cells; or (5) unidentified aspects of the culture method used in this study may promote cell differentiation [9].

Cell therapy for kidney diseases using stem cells is currently under investigation. Mesenchymal stem cell transfer has been suggested to improve the prognosis of acute kidney injury by exerting immunomodulatory effects and to benefit chronic kidney disease by suppressing angiogenesis and fibrosis [18]. The immunomodulatory effects and differentiation capacities of transferred stem cells vary depending on their tissue of origin and the donor’s age [2][19], although their specific effects for each disease type remain to be determined [2]. Current research is also exploring whether inducing stem cell differentiation and proliferation can support kidney reconstruction and be applied in regenerative medicine. Because the kidney comprises many types of cells, its reconstruction requires stem cells with broad differentiation potential [19]. While human kidney organoids have been generated from iPS cells, the process remains technically complex [19]. Moreover, obtaining the stem cells needed for therapy often requires invasive procedures, whereas isolating them from urine avoids this problem [7]. Although further work is needed, we speculate that the stem cells detected in the urinary exfoliated cell cultures from this study may possess distinctive differentiation abilities that could prove particularly beneficial for certain, as yet unidentified, kidney disease types. The ease of isolation from urine could also facilitate their application in regenerative medicine by simplifying kidney tissue reconstruction procedures. In summary, this study expands the possibilities for generating mesodermal lineages from urinary exfoliated cells, which—though requiring further investigation—may prove valuable to the field.

## 5. Limitations

This experiment was performed with a small number of patients and kidney disease types. Moreover, the development of adipocytes, osteoblasts, and chondrocytes from isolated stem cells was not monitored over time, and gene expression patterns were not analyzed at the level of individual cell clusters.

## 6. Conclusion

This study reveals an unexpected ability of urinary exfoliated cells to generate multiple mesodermal lineages without specialized induction conditions. These findings suggest that our culture approach can provide urinary stem cells with distinct differentiation abilities. Isolating and characterizing these cells will be essential to explore their potential applications in cell therapy and regenerative treatments for kidney diseases.

## Abbreviations

ACS: Acetyl-CoA synthetase
ALP: Alkaline phosphatase
ANCA: anti-neutrophil cytoplasmic antibody associated glomerulonephritis
BSA: bovine serum albumin
C/EBP: CCAAT/enhancer-binding protein
DEX: dexamethasone
DMEM/F12: Dulbecco’s modified Eagle’s medium/Hams F12
FABP4: Fatty acid-binding protein 4
FBS: fetal bovine serum
GAPDH: Glyceraldehyde-3-phosphate dehydrogenase
HSPN: Henoch-Schönlein purpura nephritis
IBMX: isobutylmethylxanthine
IgAN: IgA nephropathy
ITS: insulin, transferrin, and selenium
LIF: Leukemia inhibitory factor
LN: Lupus nephritis
LPL: Lipoprotein lipase
MEM: minimum essential medium
MN: membranous nephropathy
PBS: phosphate-buffered saline
PPAR: Peroxisome proliferator-activated receptor
RUX2: Runt-related transcription factor 2
SOX2: SRY-box transcription factor 2
SOX9: SRY-box transcription factor 9

## Author’s Contributions

N.K. performed the experiments and wrote the manuscript; T.A., T.K., and Y.M. collected the clinical data of the patients; M.A. and Y.I. critically reviewed the manuscript for important intellectual content.

All authors have read and approved the final version of the manuscript.

## Acknowledgement

The authors thank Dr. J.M. Dijkstra at Fujita Health University for proof-reading and English language editing.

